# Chronic intermittent ethanol and acute stress similarly modulate BNST CRF neuron activity via noradrenergic signaling

**DOI:** 10.1101/458349

**Authors:** Angela E Snyder, Danny G Winder, Yuval Silberman

**Affiliations:** Department of Neural and Behavioral Sciences, Penn State College of Medicine; Vanderbilt Center for Addiction Research; Vanderbilt Brain Institute; Department of Molecular Physiology & Biophysics; Vanderbilt J.F. Kennedy Center for Research on Human Development, Vanderbilt University School of Medicine

## Abstract

Relapse is a critical barrier to effective long-term treatment of alcoholism, and stress is often cited as a key trigger to relapse. Numerous studies suggest that stress-induced reinstatement to drug seeking behaviors is mediated by norepinephrine (NE) and corticotropin releasing factor (CRF) signaling interactions in the bed nucleus of the stria terminalis (BNST), a brain region critical to many behavioral and physiologic responses to stressors. Here we sought to directly examine the effects of NE on BNST CRF neuron activity and determine if these effects may be modulated by chronic intermittent EtOH (CIE) exposure or a single restraint stress. Utilizing whole-cell patch clamp electrophysiological techniques in *CRF-tomato* reporter mice, we found that NE depolarized BNST CRF neurons in naïve mice in a β-adrenergic receptor (AR) dependent mechanism. CRF neurons from CIE or stress-exposed mice had significantly elevated basal resting membrane potential compared to naïve mice. Furthermore, CIE and stress individually disrupted the ability of NE to depolarize CRF neurons, suggesting that both stress and CIE utilize β-AR signaling to modulate BNST CRF neurons. Neither stress nor CIE altered the ability of exogenous NE to inhibit evoked glutamatergic transmission onto BNST CRF neurons, a mechanism previously shown to be α-AR dependent. Altogether these findings suggest that stress and CIE interact with β-AR signaling to modulate BNST CRF neuron activity, potentially disrupting the α/β-AR balance of BNST CRF neuronal excitability. Restoration of α/β-AR balance may lead to novel therapies for the alleviation of many stress-related disorders.

## Introduction

The bed nucleus of the stria terminalis (BNST) plays a critical role in the behavioral and physiologic responses to stress (Casada and Dafny 1991; Crestani et al. 2013; Sullivan et al. 2004; Tran et al. 2012; Waddell et al. 2006). In particular, corticotropin releasing factor (CRF) signaling is thought to be an important regulator of pro-anxiety BNST-mediated responses to stress (Ciccocioppo et al. 2003; Koob 1999; Sink et al. 2013; Walker et al. 2009). Numerous studies have shown that CRF signaling is enhanced in the BNST following acute and chronic stress exposures (Choi et al. 2006; Funk et al. 2006; Janitzky et al. 2014; Tran et al. 2012). In addition, a number of studies indicate that stress or adverse external stimuli can increase norepinephrine (NE) release in the BNST (Flavin and Winder 2013; Fuentealba et al. 2000; Pacak et al. 1995; Park et al. 2012). BNST NE signaling is critical for stress-related increases in drug-seeking behaviors (Brown et al. 2011; Leri et al. 2002; Wang et al. 2001), behaviors that require CRF receptor activation (Brown et al. 2009; Erb et al. 2001; McReynolds et al. 2014). Moreover, anatomic and electrophysiologic studies indicate that noradrenergic terminals interact with CRF producing neurons in the BNST (Nobis et al. 2011; Phelix et al. 1994) suggesting stress-induced NE release in the BNST can increase CRF neuron activity to coordinate behavioral responses to stress. Furthermore, this circuitry is sensitive to modulation by chronic ethanol (EtOH) (Olive et al. 2002; Silberman et al. 2013). Together, these studies implicate NE modulation of BNST CRF neurons in anxiogenic behaviors and in stress-induced EtOH seeking behaviors. However, the mechanisms by which stress and chronic EtOH modulate BNST CRF neuron activity has not been fully elucidated.

To that end, the current study sought to directly examine the mechanism by which stress and chronic EtOH exposure alter BNST CRF neuron excitability. Our findings suggest that stress and chronic EtOH exposure enhance BNST CRF neuron activity via similar β-AR dependent mechanisms. Surprisingly, stress and chronic EtOH do not appear to alter NE-induced inhibition of glutamatergic inputs onto BNST CRF neurons, an effect previously shown to be α-AR dependent (Fetterly et al. 2016). Together, these findings indicate that stress and chronic EtOH target the activity of excitatory β-ARs on BNST CRF neurons without altering inhibitory α-AR modulation of these neurons, thereby altering the α/β-AR balance within this circuitry. These findings further suggest that maintaining α/β-AR balance in BNST CRF circuits may be an important target for novel treatments for stress-related disorders and stress-induced reinstatement to alcohol seeking behaviors.

## Methods

### Animals

Adult (>7wk old) male CRF-tomato reporter mice (Silberman et al. 2013) were used in all studies. CRF neurons were identified for electrophysiological analysis through the use of CRF-*tomato* reporter mice as previously described (Silberman et al. 2013). The CRF-Cre line utilized to produce the CRF-tomato mice in these studies has been extensively evaluated and reliably reports CRF mRNA-expressing neurons (Chen et al. 2015). All mice were housed in groups of two to five for the duration of the studies. Food and water were available *ad libitum.* All procedures were approved by the Institutional Animal Care and Use Committees at Vanderbilt University (Nashville, TN) and Penn St College of Medicine (Hershey, PA).

### Restraint stress exposure

Mice were allowed to acclimate to the test location in their home cage for one hour in a sound- and light-attenuating box (Med Associates Inc.). Mice were restrained during the light cycle (0900-1100) in restraint devices made from 50-mL Falcon conical tubes (Fisher Scientific) with several (≈15) holes in the front and rear (cap) to maintain airflow (McElligott et al. 2010). While in the restraint devices, animals were placed in individual cages inside sound- and light-attenuating boxes in the same room for 1 hour, and then returned to their home cage for 30 min prior to preparation for electrophysiology experiments.

### Chronic Intermittent Ethanol (CIE)

CIE procedures were performed as previously published (Silberman et al. 2013). Briefly, mice were given a daily injection of pyrazole (1 mmol/kg), placed in a chamber filled with volatilized ethanol for 16 hours per day, then returned to standard animal housing for 8 hours. Mice were exposed to CIE for 4 days (one CIE cycle) followed by three days solely in standard animal housing before being returned to the chamber for a second CIE cycle. On the last day of the second CIE cycle, mice were returned to the standard animal housing facility for either 4 hours or 4 days before being used for electrophysiology.

### Electrophysiology

250-300 μm-thick coronal brain slices containing the BNST (Bregma, +0.14–0.26) were prepared from adult male CRF-tomato mice as previously described (Kash et al. 2008; Nobis et al. 2011). Following acclimation, mice were briefly anesthetized with isoflurane and brains were quickly removed and submerged in ice-cold, oxygenated low-sodium sucrose dissecting solution (in mM: 194 sucrose, 20 NaCl, 4.4 KCl, 2 CaCl_2_, 1 MgCl_2_, 1.2 NaH_2_PO_4_, 10 glucose, 26 NaHCO_3_). A vibratome (Leica) was used to prepare the brain slices, which were then transferred to a holding chamber containing oxygenated ACSF at 28°C, and allowed to recover for at least one hour prior to recordings.

Whole-cell voltage-clamp recordings of AMPA receptor-mediated excitatory postsynaptic currents (EPSCs) were made at −70 mV and pharmacologically isolated by the addition of 25 μM picrotoxin (Tocris) to the standard ACSF (in mM: 124 NaCl, 4.4 KCl, 2 CaCl2, 1.2 MgSO4, 1 NaH2PO4, 10 glucose, and 26 NaHCO3). Recording electrodes for voltage-clamp experiments were filled with a cesium gluconate internal solution (in mM: 118 CsOH, 117 D-gluconic acid, 5 NaCl, 10 HEPES, 0.4 EGTA, 2 MgCl2, 5 tetraethylammonium chloride, 4 ATP, 0.3 GTP, pH 7.2–7.3, 270–290 mOsmol). A 50 msec interevent interval was used to examine paired-pulse ratio (PPR). Whole-cell current-clamp recordings were carried out at each neuron’s resting membrane potential performed with the addition of 25 μM picrotoxin and 3 mM kynurenic acid to the ACSF (to block GABA_A_ and AMPA/NMDA mediated neurotransmission, respectively) and recording electrodes were filled with a potassium gluconate internal solution (in mM: 135 K+-gluconate, 5 NaCl, 2 MgCl2, 10 HEPES, 0.6 EGTA, 4 ATP, 0.4 GTP, pH 7.2–7.3, 280–290 mOsmol). When used, the β-AR antagonist propranolol was pre-applied for at least 15 min before recordings began and remained onboard for the duration of the experiment. All electrophysiology recordings were made using Clampex 9.2 and analyzed using Clampfit 10.2-10.7 (Molecular Devices). Experiments in which the access resistance changed by >20% or were otherwise unstable were not included in the data analyses.

### Statistical analyses

Statistical analyses were performed using Microsoft Excel 2011 and GraphPad Prism 7. To determine if a drug had a significant effect we compared the experimental values to baseline values using a Student’s paired t-test. Unpaired t-tests were used to determine significant differences of a drug effect between two groups. Differences between three or more groups were assessed using one-way ANOVA followed by Fisher’s LSD post-hoc test to determine the significance of a specific group compared to stress/EtOH-naïve control mice. All values throughout the study are presented as mean ± SEM. Graphpad Prism 7 and Powerpoint 2011 were used for figure preparation.

### Reagents

All reagents used were purchased from MilliporeSigma unless otherwise noted in the text.

## Results

### Norepinephrine depolarizes BNST CRF neurons via β-AR activation

Our previous findings indicate that β-AR agonists can depolarize BNST CRF neurons and increase excitatory neuronal transmission onto BNST projection neurons in a CRF-dependent mechanism (Silberman et al. 2013). These findings suggest that NE may be a critical signaling component in regulating BNST CRF neuron excitability. Here, we sought to determine if NE may directly alter BNST CRF neuron excitability, and if this mechanism is sensitive to CIE or stress. To that end, we recorded from *tomato+* neurons in the BNST of CRF reporter mice and measured neuron resting membrane potential (RMP) in the presence of both GABA and glutamate receptor antagonists (25 μM picrotoxin and 3 mM kynurenic acid) to remove potential confounds of previously described NE modulation of BNST network activity (Flavin and Winder 2013). Bath application of 1 μM NE significantly depolarized BNST CRF neurons compared to basal RMP (5.8±1.3 mV, n=12, p<0.005, Fig 1A-B). The magnitude of NE-induced depolarization was significantly correlated with a decrease in input resistance (r^2^=0.8689, p<0.0001, Fig 1C). In some cases, BNST CRF neurons that became depolarized began to spontaneously fire action potentials (Fig 1A, 3 of 12 cells tested). To test the hypothesis that β-AR stimulation is required for NE-induced depolarization of BNST CRF neurons, we pretreated slices with 10 μM propranolol, a β-AR antagonist, for a minimum of 15 minutes prior to recording BNST CRF neuron RMP. In the presence of propranolol pretreatment, NE did not significantly alter BNST CRF neuron RMP (0.17±1.2 mV change from baseline, n=6; p>0.05 compared to baseline, Fig 1D-E). The ability of NE to depolarize BNST CRF neuron membrane potentials was significantly different in control vs propranolol treated cells (p<0.01, Fig 1F).

**Figure 1.**
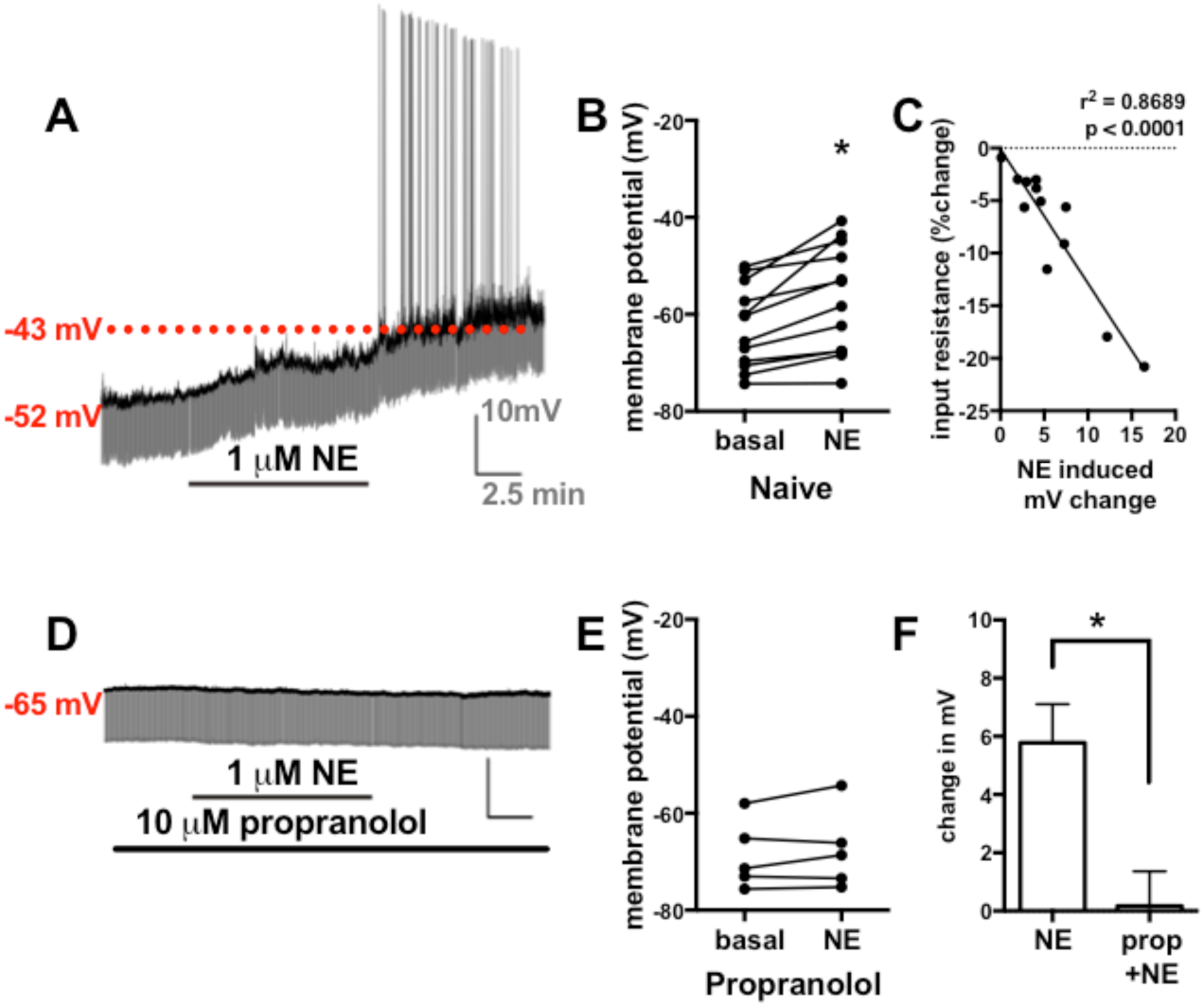
NE depolarizes BNST CRF neurons in a β-AR dependent manner. A) Example trace showing the effect of 1 μM NE on BNST CRF neuron membrane potential. Dark line indicates resting membrane potential, gray area indicates input resistance measurements. Numbers to the left indicate membrane potential (in mV) during baseline and after 10 min NE application. Solid line below trace indicates time of drug application. Scale bar: x-axis = 2.5 min; y-axis = 10 mV. B) Line graph showing the effect of NE on BNST CRF neuron membrane potential compared to basal levels on individual neurons tested. * indicates a significant difference from basal (p<0.05). C) Analysis showing that magnitude of NE-induced depolarization is significantly correlated with larger changes to input resistance. Dashed line indicates input resistance normalized to 0% change. D) Example trace showing the effect of 1 μM NE on BNST CRF neuron membrane potential in slices pretreated with the β-AR antagonist propranolol. Dark line indicates resting membrane potential, gray area indicates input resistance measurements. Solid lines below trace indicate time of drug applications. Number to the left indicates membrane potential (in mV) during baseline. Scale bar: x-axis = 2.5 min; y-axis = 10 mV. E) Line graph showing the effect of propranolol pretreatment on NE modulation of BNST CRF neuron membrane potential on individual neurons tested. F) Bar graph summarizing the difference in NE effects on change in BNST CRF neuron membrane potentials with and without propranolol pretreatment. * indicates significant difference between groups (p<0.05).

### Acute stress and short-term CIE withdrawal shift BNST CRF neuron RMP

Mice underwent CIE for two cycles and either acute withdrawal (4 hr) or extended withdrawal (4 days) prior to slice preparation. Recordings of BNST CRF neuron RMPs were performed as above. A separate cohort of mice was EtOH naïve but exposed to a 1 hr restraint stress before slice preparation as previously published (Fetterly et al. 2016). One-way ANOVA revealed significant effects of stress and CIE on BNST CRF neuron RMPs (F_(3,29)_=3.376, p<0.05, Fig 2A) compared to cells from EtOH/stress naïve mice. Fisher’s LSD post-hoc test showed a significant difference between naïve (RMP= −62.6±2.5 mV, n=12) and stress-exposed mice (−51.8±6.2, n= 5, p<0.05) and between naïve and 4 hr CIE withdrawal mice (−54.7±2.1, n= 10, p<0.05), but no significant difference between naïve and 4 day CIE withdrawal mice (−63.4±1.6, n=6, p=0.85).

**Figure 2.**
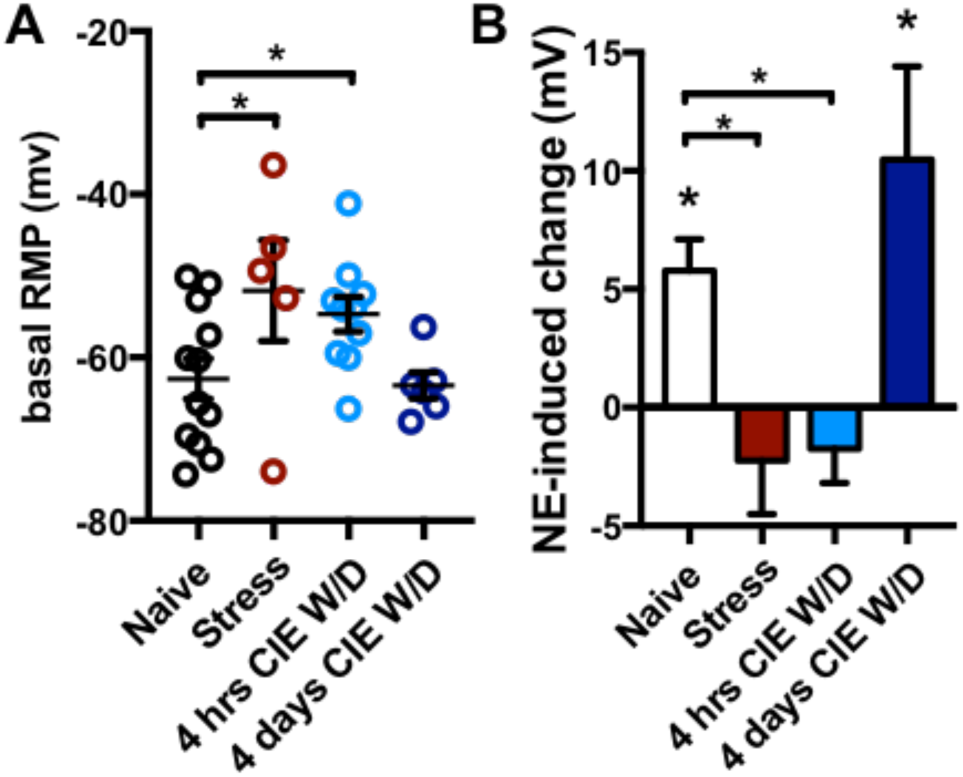
Effects of NE on BNST CRF neuron resting membrane potentials in stress and CIE exposed mice. A)Scatter plot showing basal resting membrane potentials (RMP) in EtOH/stress Naïve mice (same cells as in figure 1) vs mice exposed to 1 hr restraint stress or CIE exposed mice at 4 hr or 4 day withdrawal periods. * with brackets indicates significant difference between groups. B) Bar graph summarizing the effects of NE-induced changes to BNST CRF neuron membrane potential in naive vs restraint stress or CIE exposed mice at 4 hr or 4 day withdrawal periods. Basal RMP is normalized to 0. * with brackets indicates significant difference between groups. * without brackets indicates a significant increase from basal RMPs

### NE depolarization of BNST CRF neurons is occluded by acute stress or CIE withdrawal

We next sought to determine if CIE or stress may modulate the ability of NE to depolarize BNST CRF neurons (Fig 2B). After basal RMPs were determined (Fig 2A), 1 μM NE was bath applied for ten minutes as in Fig 1. One-way ANOVA revealed significant effects of stress and CIE on NE-induced changes to BNST CRF neuron RMPs (F_(3,26)_=6.302, p<0.005) compared to cells from EtOH/stress naïve mice. Fisher’s LSD post-hoc analysis showed a significant difference between naïve (NE-induced change = 5.8±1.3 mV, n=12) and stress mice (−0.9±2.2 mV, n= 5, p<0.05) and between naïve and 4 hr CIE withdrawal mice (−1.741 ±1.4, n= 7, p<0.05). The effects of NE on BNST CRF neuron RMPs in the stress and 4 hr CIE withdrawal group were not significantly different from baseline (p>0.05). However, NE was able to significantly alter BNST CRF neuron RMPs from baseline in the 4 day CIE withdrawal group (10.5±3.9mV, n=6, p<0.05) and this effect was not different from naïve mice (p=0.12).

### Stress and CIE do not modulate NE inhibition of glutamatergic transmission onto BNST CRF neurons

Together, the above findings indicate that NE can depolarize BNST CRF neurons via a β-AR dependent mechanism that is functionally occluded following acute stress or short-term CIE withdrawal. Previous work indicates that NE can inhibit glutamatergic transmission in the BNST via a distinct α-AR mediated mechanism (Flavin and Winder 2013; Forray et al. 1999; Fuentealba et al. 2000). Thus, we next sought to determine if stress or CIE would alter NE modulation of glutamatergic transmission onto BNST CRF neurons. After restraint stress, bath application of 1 μM NE significantly inhibited EPSC amplitude in BNST CRF neurons (−61.7 ±13.1% change from baseline, n=5, p<0.01, Fig 3) without altering PPR (14.3±10.1% change from baseline, p>0.05). The effect of NE in restraint stress exposed mice is similar to previous findings in stress-naïve animals (Fetterly et al. 2016).

**Figure 3.**
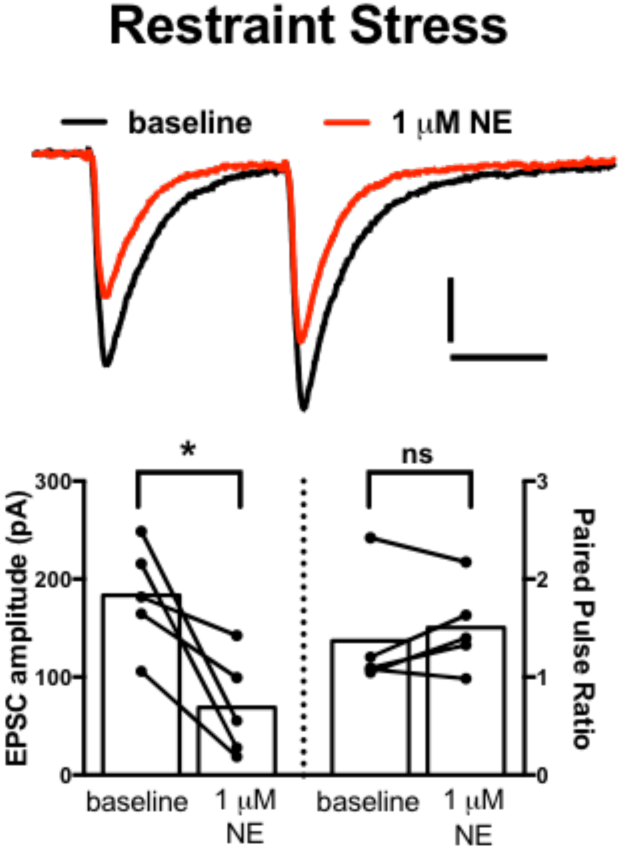
NE inhibits glutamatergic transmission in BNST-CRF neurons in stress exposed mice. Top) Example traces of the effect of 1 μM NE on electrically evoked excitatory postsynaptic currents (EPSC) in BNST cRf neurons from animals exposed to 1-hr restraint stress. Scale bar: x axis = 25 msec; y axis = 50 pA. Bottom) Before-and-after line and bar graphs summarizing the effects of NE on EPSC amplitude and paired pulse ratio. * indicates a significant difference from baseline conditions. ns= not significant.

In 4 hr CIE withdrawal mice, 1 μM NE also significantly decreased EPSC amplitude in BNST CRF neurons (−34.3±7.9% change from baseline, n=5, p<0.05, Fig 4) without altering PPR (0.1 ±4.6, p>0.05). The effect of NE on evoked EPSCs in 4 hr CIE withdrawal mice was not different from effects in stressed mice (p>0.05) and was similar to previous data in naïve mice (Fetterly 2016). Together, these findings indicate that stress and CIE exposure do not alter the ability of NE to inhibit excitatory drive onto BNST CRF neurons. As there was no difference in the 4 hr CIE withdrawal group in this experiment, we did not test NE effects at the 4 day CIE withdrawal time point.

**Figure 4.**
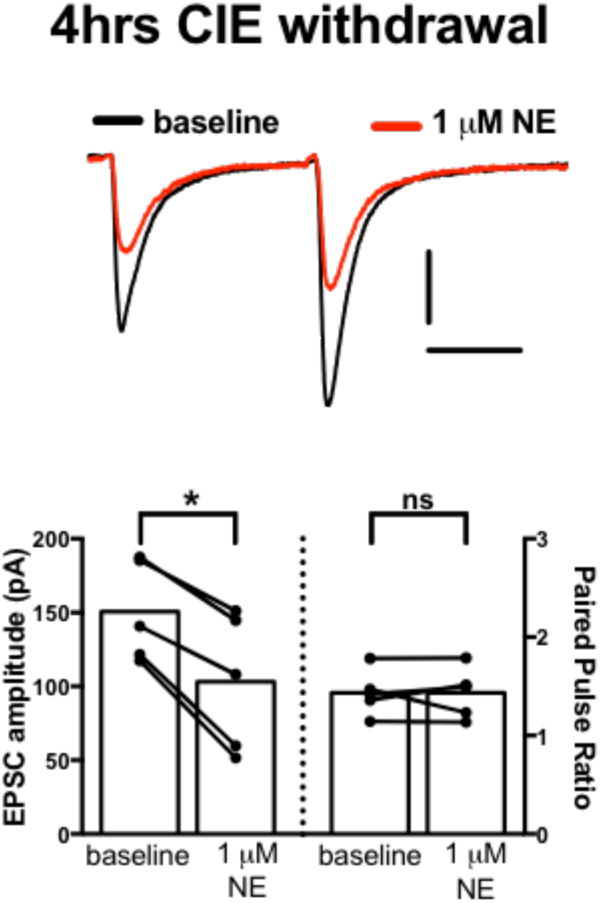
NE inhibits glutamatergic transmission in BNST-CRF neurons in CIE exposed mice. Top) Example traces of the effect of 1 μM NE on electrically evoked excitatory postsynaptic currents (EPSC) in BNST CRf neurons from animals following 4 hr withdrawal from CIE. Scale bar: x axis = 25 msec; y axis = 50 pA. Bottom) Before-and-after line and bar graphs summarizing the effects of NE on EPSC amplitude and paired pulse ratio. * indicates a significant difference from baseline conditions. ns= not significant.

## Discussion

Noradrenergic activation of BNST CRF neurons is thought to be critically involved in physiologic and behavioral responses to stress, and is a key component driving stress-induced reinstatement to drug-seeking behaviors (Brown et al. 2011; Leri et al. 2002; Wang et al. 2001). Here we provide novel, direct evidence that NE can depolarize BNST CRF neurons via β-AR stimulation. Furthermore, BNST CRF neurons had significantly more positive RMPs following restraint stress or CIE exposure. In addition, the depolarizing effects of exogenous NE application were functionally occluded by restraint stress or CIE exposure. While previous work indicated that NE could also inhibit glutamatergic transmission onto BNST CRF neurons via an α-AR mediated mechanism (Fetterly et al. 2016), this effect of NE is not modulated by exposure to either restraint stress or CIE. Together, these findings suggest NE can alter BNST CRF neuron excitability via modulation of two mechanisms: 1) direct depolarization of BNST CRF neurons via a β-AR dependent mechanism that is stress and ethanol sensitive and 2) an α-AR dependent mechanism that inhibits glutamatergic transmission onto BNST CRF neurons that is not sensitive to stress or CIE exposure. Therefore, NE modulation of BNST CRF neurons appears to occur via balanced α/β-AR signaling, and stress and CIE shift this balance by preferentially targeting β-ARs.

A number of previous studies suggested that a variety of stressors and drugs of abuse enhance neuronal activity in the BNST in general as well as enhance the activity of BNST CRF neurons specifically (Casada and Dafny 1991; Funk et al. 2006; Janitzky et al. 2014; Tran et al. 2012; Walker et al. 2009). Previous data also indicate that CRF signaling may be an important factor in overall BNST excitability and that BNST CRF neurons are sensitive to β-AR stimulation. For instance, the β-AR agonist isoproterenol can increase glutamatergic transmission in the BNST via a CRF receptor dependent mechanism (Nobis et al. 2011) and can depolarize BNST CRF neurons (Silberman et al. 2013). CRF has also been shown to enhance glutamatergic transmission onto BNST projection neurons, an effect that is mimicked by CIE (Silberman et al. 2013). Together, these findings suggest that stress and CIE mediated β-AR activation can stimulate the release of endogenous CRF in the BNST from local CRF producing neurons to enhance BNST excitability. In support of this hypothesis, the current findings show that NE can directly depolarize BNST CRF neurons, in some cases resulting in spontaneous action potential firing. This effect is dependent upon β-AR stimulation and is functionally occluded in restraint stress and chronic ethanol exposed mice. BNST CRF neurons’ resting membrane potentials are more positive in stress and CIE withdrawal mice compared to naïve animals, suggesting endogenous β-AR signaling may have already occurred. Overall these data provide direct evidence for neurocircuit function previously hypothesized from studies of stress and chronic ethanol activation of the BNST circuits.

An alternative hypothesis for NE-induced enhancement of BNST CRF neuron activity could be that NE may increase excitatory glutamatergic transmission onto BNST CRF neurons. However, recent evidence suggests that this is not likely to be the case as NE inhibits glutamatergic transmission onto BNST CRF neurons (Fetterly et al. 2016). Analysis with AR-selective agonists/antagonists suggests that NE inhibits glutamatergic transmission onto BNST CRF neurons via a predominantly α-AR mediated mechanism. This is an important consideration as α1-AR stimulation results in a long-term depression (LTD) of excitatory transmission in the BNST (McElligott and Winder 2008), suggesting an important role for these receptors in regulating overall BNST excitation. Intriguingly, NE inhibition of glutamatergic transmission in BNST CRF neurons is maintained following CIE or stress exposure in this study. The ability of α-ARs to maintain NE sensitivity following a stress or CIE exposure may be a mechanism by which BNST CRF neurons return to homeostatic activity in response to a prolonged β-AR activation. While the data here examine an acute stressor, α1-AR LTD of glutamatergic transmission in the BNST is disrupted by chronic/repeated restraint stress (McElligott et al. 2010), suggesting that pathology of α1-AR mediated signaling may be an important factor in the development of stress-related disorders via loss of the ability of CRF neurons to return to a homeostatic state. It will be important in future studies to further examine the mechanisms by which α1-AR signaling in BNST CRF neurons may be disrupted by chronic stress as an avenue for the development of novel treatments for these disorders.

α2-ARs are also involved in regulating synaptic transmission in the BNST (Shields et al. 2009), particularly inputs from the parabrachial nucleus to CRF neurons (Fetterly et al. 2016; Flavin et al. 2014). While the role of parabrachial inputs to the BNST are still being elucidated, α2-AR agonists can inhibit the ability of stress to enhance cFos activity in BNST CRF neurons (Fetterly et al. 2016). This finding indicates multiple α-AR mediated recovery mechanisms may be targeted in the BNST CRF circuit. Overall the findings here suggest that the ability of acute stress, CIE withdrawal, and NE to enhance BNST CRF neuron activity is likely due to stimulation of β-ARs on BNST CRF neurons and not via α-AR modulation of excitatory transmission onto BNST CRF neurons.

In conclusion, these findings provide novel direct electrophysiologic evidence to support the hypothesis that stress and CIE enhance BNST excitability via β-AR mediated stimulation of BNST CRF neurons. Previous work combined with the findings here indicate that α- and β-AR work in conjunction, via distinct mechanisms and potentially distinct circuits, to regulate overall BNST excitability and BNST CRF neuron activity in particular. Disrupting the balance of α- and β-ARs in the BNST may be a critical component of stress and EtOH induced behavioral changes. Recent evidence also suggests that competing pro-stress and anti-stress circuits are present in the BNST(Daniel and Rainnie 2016), the balanced activity of which drives appropriate behavioral responses to stress. While the data here indicate that CIE modulation of β-AR signaling can return to normal levels after 4 days of withdrawal, it will be important in future studies to determine if stress, CIE, or their combined exposures will lead to long-lasting changes in α/β-AR balance of BNST CRF neuron excitability, or differentially alter the activity of competing pro- and anti-anxiety circuits in the BNST.

## Acknowledgements

This work was supported by National Institutes of Health Grants R01DA042475 (DGW), R01DA042475S1 (DGW), and R00AA022937 (YS).

## References

Brown ZJ, Nobrega JN, Erb S. 2011. Central injections of noradrenaline induce reinstatement of cocaine seeking and increase c-fos mrna expression in the extended amygdala. Behav Brain Res. 217(2):472–476.

Brown ZJ, Tribe E, D’souza NA, Erb S. 2009. Interaction between noradrenaline and corticotrophin-releasing factor in the reinstatement of cocaine seeking in the rat. Psychopharmacology. 203(1):121–130.

Casada JH, Dafny N. 1991. Restraint and stimulation of bed nucleus of the stria terminalis produce similar stress-like behaviors. Brain research bulletin. 27(2):207–212.

Chen Y, Molet J, Gunn BG, Ressler K, Baram TZ. 2015. Diversity of reporter expression patterns in transgenic mouse lines targeting corticotropin-releasing hormone-expressing neurons. Endocrinology. 156(12):4769–4780.

Choi DC, Nguyen MM, Tamashiro KL, Ma LY, Sakai RR, Herman JP. 2006. Chronic social stress in the visible burrow system modulates stress-related gene expression in the bed nucleus of the stria terminalis. Physiology & behavior. 89(3):301–310.

Ciccocioppo R, Fedeli A, Economidou D, Policani F, Weiss F, Massi M. 2003. The bed nucleus is a neuroanatomical substrate for the anorectic effect of corticotropin-releasing factor and for its reversal by nociceptin/orphanin fq. The Journal of Neuroscience. 23(28):9445–9451.

Crestani CC, Alves FHF, Gomes FV, Resstel LBM, Correa FMA, Herman JP. 2013. Mechanisms in the bed nucleus of the stria terminalis involved in control of autonomic and neuroendocrine functions: A review. Current Neuropharmacology. 11(2): 141–159.

Daniel SE, Rainnie DG. 2016. Stress modulation of opposing circuits in the bed nucleus of the stria terminalis. Neuropsychopharmacology: official publication of the American College of Neuropsychopharmacology. 41(1):103–125.

Erb S, Shaham Y, Stewart J. 2001. Stress-induced relapse to drug seeking in the rat: Role of the bed nucleus of the stria terminalis and amygdala. Stress. 4(4):289–303.

Fetterly T, Awad EK, Silberman Y, Winder DG. 2016. Control of bnst crf neurons by norepinephrine and stress [abstract]. Society for Neuroscience; San Diego, CA.

Flavin SA, Matthews RT, Wang Q, Muly EC, Winder DG. 2014. Alpha(2a)-adrenergic receptors filter parabrachial inputs to the bed nucleus of the stria terminalis. The Journal of neuroscience: the official journal of the Society for Neuroscience. 34(28):9319–9331.

Flavin SA, Winder DG. 2013. Noradrenergic control of the bed nucleus of the stria terminalis in stress and reward. Neuropharmacology. 70:324–330.

Forray MI, Bustos G, Gysling K. 1999. Noradrenaline inhibits glutamate release in the rat bed nucleus of the stria terminalis: In vivo microdialysis studies. J Neurosci Res. 55(3):311–320.

Fuentealba JA, Forray MI, Gysling K. 2000. Chronic morphine treatment and withdrawal increase extracellular levels of norepinephrine in the rat bed nucleus of the stria terminalis. J Neurochem. 75(2):741–748.

Funk D, Li Z, Lê AD. 2006. Effects of environmental and pharmacological stressors on c-fos and corticotropin-releasing factor mrna in rat brain: Relationship to the reinstatement of alcohol seeking. Neuroscience. 138(1):235–243.

Janitzky K, Peine A, Kröber A, Yanagawa Y, Schwegler H, Roskoden T. 2014. Increased crf mrna expression in the sexually dimorphic bnst of male but not female gad67 mice and tmt predator odor stress effects upon spatial memory retrieval. Behavioural Brain Research. 272(0):141–149.

Kash TL, Nobis WP, Matthews RT, Winder DG. 2008. Dopamine enhances fast excitatory synaptic transmission in the extended amygdala by a crf-r1-dependent process. The Journal of neuroscience: the official journal of the Society for Neuroscience. 28(51):13856–13865.

Koob GF. 1999. Corticotropin-releasing factor, norepinephrine, and stress. Biological Psychiatry. 46(9):1167–1180.

Leri F, Flores J, Rodaros D, Stewart J. 2002. Blockade of stress-induced but not cocaine-induced reinstatement by infusion of noradrenergic antagonists into the bed nucleus of the stria terminalis or the central nucleus of the amygdala. The Journal of neuroscience: the official journal of the Society for Neuroscience. 22(13):5713–5718.

McElligott ZA, Klug JR, Nobis WP, Patel S, Grueter BA, Kash TL, Winder DG. 2010. Distinct forms of gq-receptor-dependent plasticity of excitatory transmission in the bnst are differentially affected by stress. Proceedings of the National Academy of Sciences of the United States of America. 107(5):2271–2276.

McElligott ZA, Winder DG. 2008. Alpha1-adrenergic receptor-induced heterosynaptic long-term depression in the bed nucleus of the stria terminalis is disrupted in mouse models of affective disorders. Neuropsychopharmacology: official publication of the American College of Neuropsychopharmacology. 33(10):2313–2323.

McReynolds JR, Vranjkovic O, Thao M, Baker DA, Makky K, Lim Y, Mantsch JR. 2014. Beta-2 adrenergic receptors mediate stress-evoked reinstatement of cocaine-induced conditioned place preference and increases in crf mrna in the bed nucleus of the stria terminalis in mice. Psychopharmacology (Berl). 231(20):3953–3963.

Nobis WP, Kash TL, Silberman Y, Winder DG. 2011. Beta-adrenergic receptors enhance excitatory transmission in the bed nucleus of the stria terminalis through a corticotrophin-releasing factor receptor-dependent and cocaine-regulated mechanism. Biol Psychiatry. 69(11):1083–1090.

Olive MF, Koenig HN, Nannini MA, Hodge CW. 2002. Elevated extracellular crf levels in the bed nucleus of the stria terminalis during ethanol withdrawal and reduction by subsequent ethanol intake. Pharmacol Biochem Behav. 72(1-2):213–220.

Pacak K, McCarty R, Palkovits M, Kopin IJ, Goldstein DS. 1995. Effects of immobilization on in vivo release of norepinephrine in the bed nucleus of the stria terminalis in conscious rats. Brain research. 688(1-2):242–246.

Park J, Wheeler RA, Fontillas K, Keithley RB, Carelli RM, Wightman RM. 2012. Catecholamines in the bed nucleus of the stria terminalis reciprocally respond to reward and aversion. Biol Psychiatry. 71(4):327–334.

Phelix CF, Liposits Z, Paull WK. 1994. Catecholamine-crf synaptic interaction in a septal bed nucleus: Afferents of neurons in the bed nucleus of the stria terminalis. Brain research bulletin. 33(1):109–119.

Shields AD, Wang Q, Winder DG. 2009. Alpha2a-adrenergic receptors heterosynaptically regulate glutamatergic transmission in the bed nucleus of the stria terminalis. Neuroscience. 163(1):339–351.

Silberman Y, Matthews RT, Winder DG. 2013. A corticotropin releasing factor pathway for ethanol regulation of the ventral tegmental area in the bed nucleus of the stria terminalis. The Journal of neuroscience: the official journal of the Society for Neuroscience. 33(3):950–960.

Sink KS, Walker DL, Freeman SM, Flandreau EI, Ressler KJ, Davis M. 2013. Effects of continuously enhanced corticotropin releasing factor expression within the bed nucleus of the stria terminalis on conditioned and unconditioned anxiety. Mol Psychiatry. 18(3):308–319.

Sullivan GM, Apergis J, Bush DE, Johnson LR, Hou M, Ledoux JE. 2004. Lesions in the bed nucleus of the stria terminalis disrupt corticosterone and freezing responses elicited by a contextual but not by a specific cue-conditioned fear stimulus. Neuroscience. 128(1):7–14.

Tran L, Wiskur B, Greenwood-Van Meerveld B. 2012. The role of the anteriolateral bed nucleus of the stria terminalis in stress-induced nociception.

Waddell J, Morris RW, Bouton ME. 2006. Effects of bed nucleus of the stria terminalis lesions on conditioned anxiety: Aversive conditioning with long-duration conditional stimuli and reinstatement of extinguished fear. Behavioral neuroscience. 120(2):324–336.

Walker DL, Miles LA, Davis M. 2009. Selective participation of the bed nucleus of the stria terminalis and crf in sustained anxiety-like versus phasic fear-like responses. Progress in Neuro-Psychopharmacology and Biological Psychiatry. 33(8):1291–1308.

Wang X, Cen X, Lu L. 2001. Noradrenaline in the bed nucleus of the stria terminalis is critical for stress-induced reactivation of morphine-conditioned place preference in rats. European journal of pharmacology. 432(2-3):153–161.

